# Electric-field-driven microfluidics for rapid CRISPR-based diagnostics and its application to detection of SARS-CoV-2

**DOI:** 10.1101/2020.05.21.109637

**Authors:** Ashwin Ramachandran, Diego A. Huyke, Eesha Sharma, Malaya K. Sahoo, Niaz Banaei, Benjamin A. Pinsky, Juan G. Santiago

## Abstract

The rapid spread of COVID-19 across the world has revealed major gaps in our ability to respond to new virulent pathogens. Rapid, accurate, and easily configurable molecular diagnostic tests are imperative to prevent global spread of new diseases. CRISPR-based diagnostic approaches are proving to be useful as field-deployable solutions. In a basic form of this assay, the CRISPR-Cas12 enzyme complexes with a synthetic guide RNA (gRNA). This complex is activated when it highly specifically binds to target DNA, and the activated complex non-specifically cleaves single-stranded DNA reporter probes labeled with a fluorophore-quencher pair. We recently discovered that electric field gradients can be used to control and accelerate this CRISPR assay by co-focusing Cas12-gRNA, reporters, and target. We achieve an appropriate electric field gradient using a selective ionic focusing technique known as isotachophoresis (ITP) implemented on a microfluidic chip. Unlike previous CRISPR diagnostic assays, we also use ITP for automated purification of target RNA from raw nasopharyngeal swab sample. We here combine this ITP purification with loop-mediated isothermal amplification, and the ITP-enhanced CRISPR assay to achieve detection of SARS-CoV-2 RNA (from raw sample to result) in 30 min for both contrived and clinical nasopharyngeal swab samples. This electric field control enables a new modality for a suite of microfluidic CRISPR-based diagnostic assays.

**Significance statement:** Rapid, early-stage screening is especially crucial during pandemics for early identification of infected patients and control of disease spread. CRISPR biology offers new methods for rapid and accurate pathogen detection. Despite their versatility and specificity, existing CRISPR-diagnostic methods suffer from the requirements of up-front nucleic acid extraction, large reagent volumes, and several manual steps—factors which prolong the process and impede use in low resource settings. We here combine on-chip electric-field control in combination with CRIPSR biology to directly address these limitations of current CRISPR-diagnostic methods. We apply our method to the rapid detection of SARS-CoV-2 RNA in clinical samples. Our method takes 30 min from raw sample to result, a significant improvement over existing diagnostic methods for COVID-19.

---

Infectious diseases such as COVID-19 are a persistent global threat. Early-stage screening and rapid identification of infected patients are important during pandemics to treat the infected and to control disease spread. The frontline diagnostic tool for COVID-19 has been RT-qPCR (reverse transcription – quantitative polymerase chain reaction), and protocols for this have been developed and published by the World Health Organization (WHO)(1) and the Centers for Disease Control and Prevention (CDC)(2). While these tests are highly specific and sensitive, they are laborious, time-consuming, and are designed for large, centralized diagnostic labs.

CRISPR-based diagnostic methods, including diagnostic assays for the COVID-19 pandemic (3, 4), have sparked great interest due to their versatility, sensitivity, and high specificity. Despite advantages, several factors have impeded highly automated CRISPR-based detection methods. For example, the CRISPR-Cas12a-based method for SARS-CoV-2 developed by Broughton et al. (3) in March 2020 requires upfront nucleic acid extraction and sample purification with a traditional adsorption/desorption column for purification which typically takes up to 1 hour. Moreover, the latter work carried out CRISPR enzymatic reactions in Eppendorf tubes and explored both colorimetric (using a lateral flow strip) and fluorescence readouts for target detection. Such protocols are not easily amenable to automation, consume significant reagent volume, and require 1 h or greater to complete. The consumption of reagents is important as Joung et al. (4) reported supply chain constrains in procuring RPA (recombinase polymerase amplification) reagents for a CRISPR-Cas13-based test which they initially developed for SARS-CoV-2 in February 2020. This limitation made them redesign their assay to one based on CRISPR-Cas12b and loop-mediated isothermal amplification (LAMP) in May 2020.

Microfluidics offers important alternate strategies to accelerate biochemical reactions (5), multiplex (6), and automate CRISPR-diagnostics. We here develop a novel electric-field-enhanced microfluidic method that is broadly applicable to the field of CRISPR-diagnostics. To this end, we use an electrokinetic microfluidic technique called isotachophoresis (ITP). ITP uses a two-buffer system which consists of a high-mobility leading electrolyte (LE) and a low-mobility trailing electrolyte (TE) buffer. On application of an electric field, sample ions with effective mobilities bracketed by the LE and TE ions selectively focus within an order 10 μm zone at the LE-to-TE interface. This focusing can preconcentrate, purify, mix, and accelerate reactions among sample and reagents species. ITP has been used to rapidly extract nucleic acids from a range of biological samples such as urine (7), blood plasma (8), and cell lysates (9), and to accelerate DNA and RNA hybridization reactions (10).

In this work, we combine microfluidics and on-chip electric field control to achieve two critical steps. First, we automatically extract nucleic acids from raw biological sample, here, nasopharyngeal swab samples from COVID-19 patients and healthy controls. Second, we use electric fields to control and effect rapid CRISPR-Cas12 enzymatic activity upon target nucleic acid recognition. The latter is achieved using a tailored on-chip ITP process to co-focus Cas12-gRNA, reporters, and target nucleic acids (Figs. S1 and S2). This creates simultaneous mixing, preconcentration, and acceleration of enzymatic reactions. Our microfluidic method consumes minimal volume of reagents on-chip and is amenable to automation. We apply our method to detection of SARS-CoV-2 in an assay which takes 30 min from raw sample to result. We demonstrate this on clinical samples, including COVID-19 positive patient samples and healthy controls. The method is both a new modality for CRISPR diagnostics and to our knowledge the fastest CRISPR-based detection of SARS-CoV-2 from raw samples with clinically relevant specificity and sensitivity.

## Results and Discussion

### Microfluidic ITP-CRISPR-based protocol for rapid SARS-CoV-2 detection from raw nasopharyngeal swab samples

Based on these principles, we developed and optimized a microfluidic protocol to rapidly detect SARS-CoV-2 viral RNA in less than 30 min starting from raw nasopharyngeal (NP) swab samples in viral transport medium (Fig. 1a). After a 2 min pre-incubation step (at 62°C) of raw NP sample with lysis buffer, we leverage on-chip ITP to rapidly extract total nucleic acids (both host DNA and any viral RNA) from the lysed sample in 3 min (Fig. 1b). Next, RT-LAMP isothermal amplification (20 min) at 62°C is performed on the ITP extract (off-chip) using a water bath, targeting the viral N and E genes and human RNase P genes in separate reactions. In the last step of our protocol, we use ITP to perform rapid (<5 min), on-chip CRISPR-Cas12-based enzymatic reactions. ITP enables simultaneous target DNA recognition by Cas12-gRNA and the resulting target-activated cleavage of ssDNA reporters. This step is carried out at room temperature and a fluorescence readout is used to detect the presence of pre-amplified nucleic acids (Fig. 1b). LAMP primers and guide RNAs for SARS-CoV-2 detection were originally published and validated by Broughton et al (3).

### ITP pre-concentrates and co-focuses Cas12-gRNA, target DNA, and reporter ssDNA into a ~100 pL reaction volume

To test our hypothesis that Cas12-gRNA can be controlled and co-focused with other reagents using an electric field gradient (in a device with no moving parts), we directly visualized the electrokinetic transport of Cas12-gRNA complex and reporter ssDNA in ITP. In these experiments, we used a Cy5-tagged (red) guide RNA and a green-fluorescent ssDNA reporter (Table S1) and we imaged the electromigration of the ITP peak near the LE-TE interface (Fig. 1c, Supplementary Video 1). The reporter ssDNA and Cas12-gRNA complex were found to electro-migrate with the same velocity as indicated by the slope (d*x*/d*t*) of the red and green fluorescence intensity fields (Fig. 1c). Further, a significant overlap of Cas12-gRNA and reporter ssDNA intensity profiles was observed experimentally (inset of Fig. 1c), which indicates these molecules co-focus in ITP— enabling reaction. Additionally, the unlabeled target DNA also co-focuses in the ITP peak since its mobility is bracketed by the LE and TE. We estimate that the concentration of Cas12-gRNA complex, target DNA and ssDNA reporters increase within ~100 pL of the ITP peak region by order ~1,000-fold compared to the initial concentrations (Fig. S3). The up-concentration of molecules within a tiny volume achieved using ITP dramatically speeds up the diffusion-limited enzymatic kinetics (11) of CRISPR-based detection assays.

### ITP-CRISPR-based detection assay for rapid detection of N and E genes of viral RNA and human RNase P gene

We next developed a novel protocol for the detection of RT-LAMP-amplified cDNA of SARS-CoV-2 viral RNA using ITP-mediated CRISPR-Cas12 DNA detection. Upon CRISPR-Cas12 binding to the target cDNA of SARS-CoV-2 viral RNA targets, Cas12 promiscuously cleaves single stranded DNA (11). The activated Cas12 cleaves reporter ssDNA probes labeled with a fluorophore-quencher pair resulting in unquenching of the fluorophore and an increase in observed fluorescence. Thus, a positive detection occurs when the fluorescence of the ITP peak rapidly increases with time (and, above a threshold), while the result is negative when there is minimal change in fluorescence (Fig. 1d and 2b, Supplementary Videos 2 and 3). We also evaluated the analytical limit of detection (LOD) of the ITP-CRISPR method (Fig. 2a). For this set of experiments, contrived samples were used which consisted of viral RNA spiked into pooled nucleic acid extracts from negative clinical nasopharyngeal swab samples. Fluorescence intensity of the moving ITP peak was measured for the N, E, and RNase P genes independently (Fig. 2b and Figs. S4, S5, and S6), and the signal at 5 min was used as the endpoint readout. The LOD of the ITP-CRISPR method was found to be 10 copies per microliter reaction, which is the same as the very recent CRISPR-based assay (3). Further, in the case of positive detection, a fluorescence signal above the threshold value was observed in < 3 min (Fig. S7). These results are in contrast to the 1 copy per microliter reaction LOD for the 2-hour qPCR method (Fig. 2e) (1). Lastly, we verified that microfluidic ITP-CRISPR detection and the typical CRISPR-based (3) approaches gave the same positive/negative result when tested with the same LAMP pre-amplified DNA (Fig. S8).

### ITP enables rapid extraction of total nucleic acids from raw nasopharyngeal swab samples

We also demonstrated on-chip ITP extraction of total nucleic acids from raw clinical positive and negative nasopharyngeal swab samples (Figs. 1b, 2a and 2d). To validate our extraction method, we performed qPCR for the E gene and RNase P control (Fig. 2e). We observed that ITP-extracted nucleic acids showed E gene amplification on positive samples, while the RNase P reaction amplified across all patient samples. These results suggest that ITP-extracted nucleic acids are compatible with downstream amplification methods.

### Demonstration of the 30-min protocol on clinical samples

Next, we combined RT-LAMP with ITP nucleic acid extraction and ITP-CRISPR detection methods and performed the complete 30-min assay (raw sample to result) on eight clinical samples (Fig. S9). A detectable fluorescence signal for the E gene and N gene targets was observed in three out of the four positive samples, while none of the negative samples exhibited a measurable signal (Figs. 2f and 2g). RNase P gene controls showed consistent positive signal in all patient samples. Using qPCR (Fig. 2e), we verified that the single positive undetected sample (Covid19-P4) was below the LOD of our ITP-CRISPR assay. A qualitative comparison of our method with other CRIPSR-based and qPCR-based detection approaches is provided in Table 1.

In summary, we developed an electrokinetic microfluidic method broadly applicable to CRISPR-based diagnostics. Our method involves ITP-based nucleic acid extraction from raw sample, isothermal reverse transcription and amplification, and then a novel CRISPR assay enhanced by ITP with a total assay time of 30 min (from raw sample to result). We applied our method to the rapid detection of SARS-COV-2 RNA from clinical nasopharyngeal swab samples. The ITP-CRISPR-based method is easily reconfigurable to the detection of novel pathogens by simply redesigning the pre-amplification primers and guide RNAs, without involving any changes to the microfluidic chip design, buffers or hardware. We hypothesize that integration of all assay steps on a single microfluidic platform (Fig. S12) can enable the development of an automated microfluidic device which achieves rapid ITP-CRISPR-based tests applicable at the point of care.

**Figure 1.**
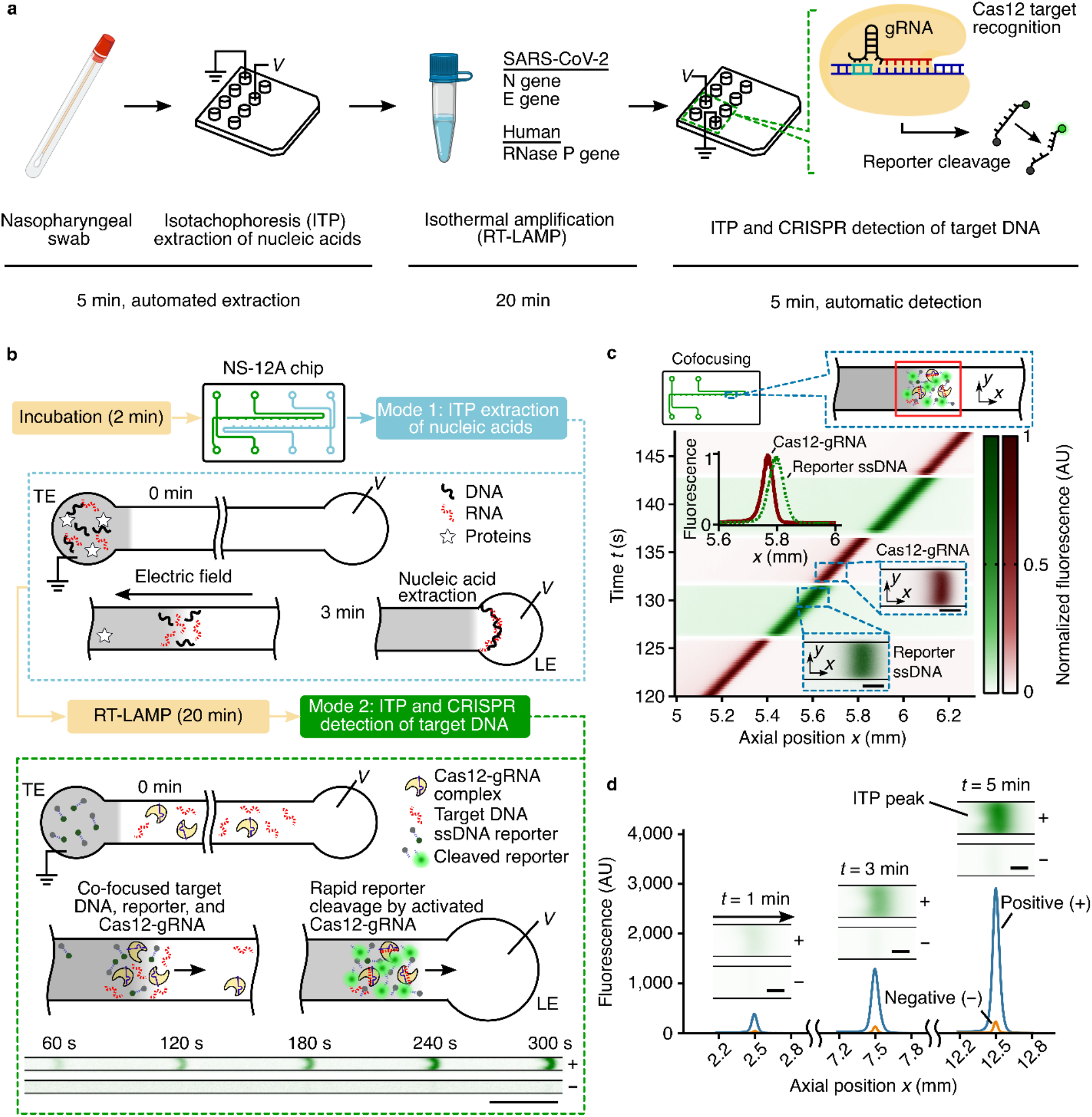
An electric-field-mediated microfluidic assay for SARS-CoV-2 RNA detection using ITP and CRISPR-Cas12. **(a)** Schematic of SARS-CoV-2 detection workflow from sample to result. Microfluidic isotachophoresis (ITP) is used to extract nucleic acids from raw nasopharyngeal sample, followed by LAMP pre-amplification and on-chip ITP-CRISPR fluorescent detection of N, E and RNase P genes. CRISPR activation by the presence of target cDNA of SARS-CoV-2 results in non-specific ssDNA cleavage and unquenching of a reporter DNA labeled with a fluorophore and quencher. **(b)** Assay working principle. A single microfluidic chip with two channels is used for ITP extraction of nucleic acids (Mode 1) and ITP-CRISPR detection (Mode 2). In Mode 1 (within dotted blue rectangle), on application of an electric field, nucleic acids with electrophoretic mobility bracketed by the leading (LE) and trailing (TE) electrolyte ions selectively focus within the electro-migrating LE-TE interface, leaving behind impurities. Following RT-LAMP of ITP-extracted nucleic acids, in Mode 2 (within the green dashed rectangle), ITP is used to effect target DNA detection using CRISPR-Cas12 enzyme assay. A positive sample shows a strong fluorescent signal compared to the negative control. Scale bar: 0.5 mm. **(c)** Electric field control of Cas12-gRNA and nucleic acids. Experimental visualization of the moving ITP interface in Mode 2 using a fluorescently tagged gRNA (red) and ssDNA reporter (green). Spatiotemporal intensity plots of the green and red fluorescence emission show that Cas12-gRNA complexes and nucleic acids electro-migrate and co-focus in a ~100 pL ITP interface volume. Top inset shows ssDNA fluorescence intensity profile (green) at 135 s and comparison with the Cas12-gRNA profile (red). Inset scale bar: 50 μm. **(d)** Example quantitative measurements of on-chip fluorescence detection from cleaving of quencher/fluorophore ssDNA reporter by ITP-focused, activated Cas12-gRNA complex. Raw on-chip fluorescence signal versus axial location for ITP-CRISPR detection of E gene of SARS-COV-2 positive and negative controls in Mode 2. Insets show instantaneous epifluorescence microscopy images of the moving ITP interface. Scale bar: 50 μm

**Figure 2.**
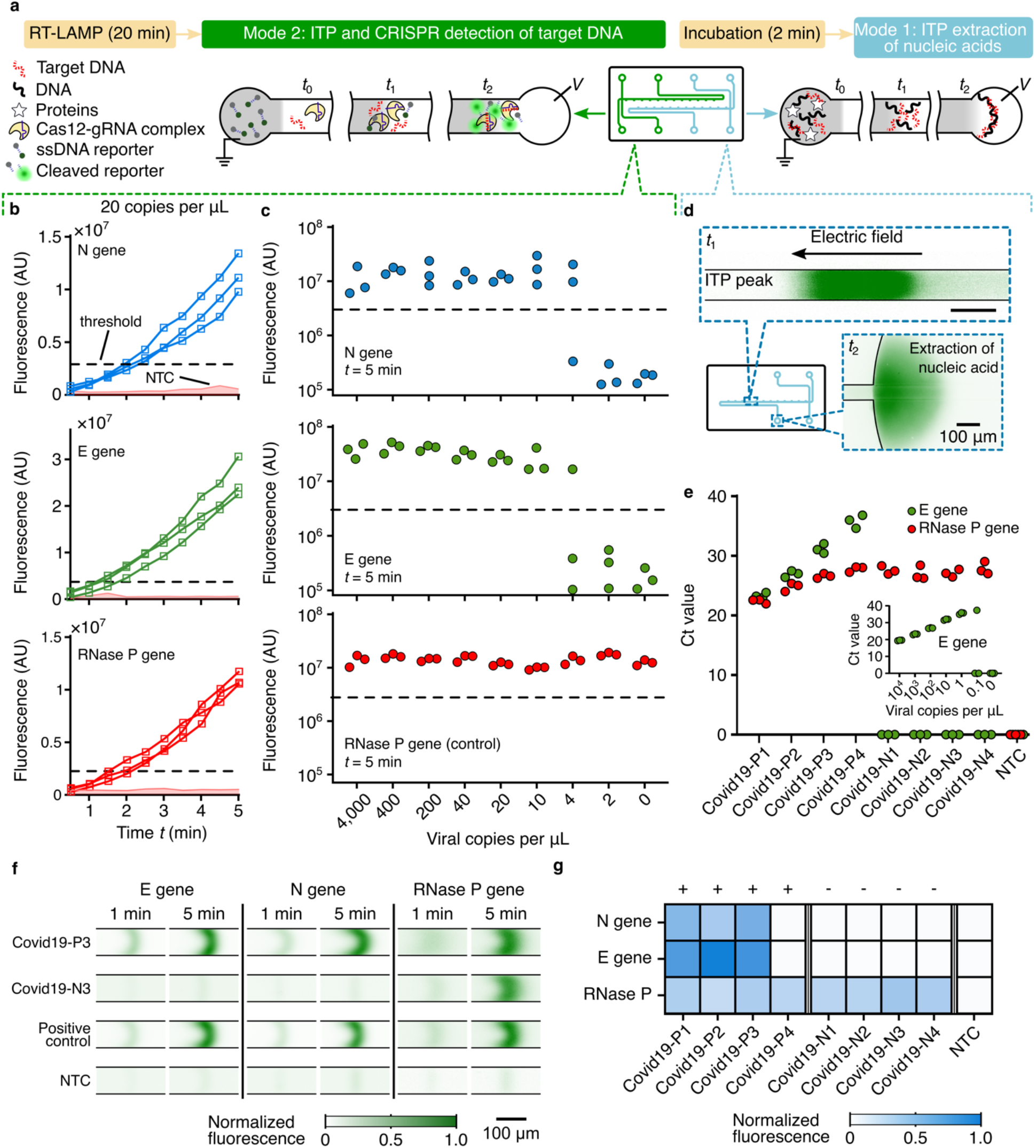
Demonstration of the assay using both contrived and clinical nasopharyngeal (NP) swab samples. **(a)** Schematic of ITP extraction and ITP-CRIPSR detection operational modes. A 2 min pre-incubation at 62°C in lysis buffer is performed prior to ITP extraction (Mode 1). 20 min LAMP at 62°C is performed prior to ITP-CRISPR detection (Mode 2). **(b)** Monitoring of fluorescence signal for contrived samples (Mode 2). Fluorescence signal for LAMP amplicons of N gene, E gene, and RNase P targets versus time for a contrived sample containing pooled nucleic acid extract from negative clinical NP swabs spiked with 20 viral genomes per microliter of reaction (*n* = 3). Shaded region represents signal from no template control (NTC) (*n* = 10). **(c)** Analytical LOD of ITP-CRISPR method. Fluorescence readout at 5 min of ITP-CRISPR assay (Mode 2). Synthetic SARS-CoV-2 RNA controls were spiked into pooled negative clinical NP swab extracts before LAMP. **(d)** Experimental images of on-chip labeled DNA and RNA focused (green) within the ITP peak during nucleic acid extraction from clinical NP sample (Mode 1). 10 μL of NP swab sample is used as input. Nucleic acids are transferred into the LE reservoir. **(e)** RT-qPCR of E gene and RNase P gene on ITP-extracted nucleic acids from clinical NP samples. Inset shows RT-qPCR standard curve for the E gene. **(f)** Fluorescence visualization of ITP peak during ITP-CRISPR detection. ssDNA reporters including quencher/fluorophore are cleaved by Cas12-gRNA on recognition of target DNA, resulting in an increased fluorescence. **(g)** Results of the complete 30-min assay on clinical nasopharyngeal swab samples. Four positive and four negative swab samples were tested. One of the positive samples (Covid19-P4) was verified to be below the 10 copies per microliter LOD of our assay. LOD: limit of detection; NTC: No template control.

**Table 1.**
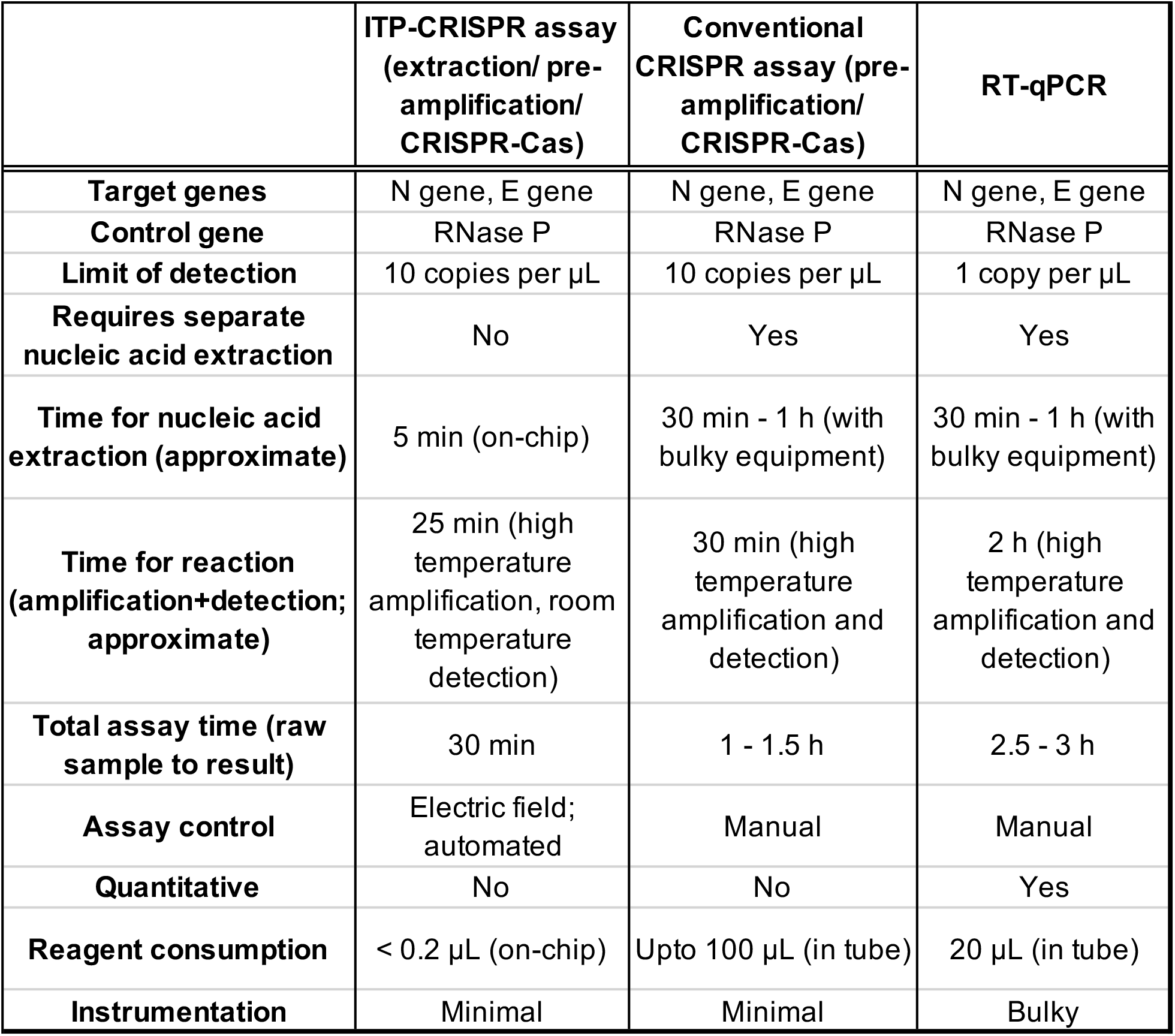
Comparison of ITP-CRISPR-based detection with the conventional CRISPR-based (3) and qPCR assays (1, 2).

## Methods

### Nucleic acid preparation

Synthetic ssRNA control for SARS-CoV-2 variant (GenBank ID: MT007544.1) was obtained (Twist Biosciences) at a concentration of one million copies per microliter. The ssRNA control sequences were generated by the transcription of six non-overlapping 5kb gene fragments of SARS-CoV-2, providing greater than 99.9% coverage of the viral genome. For analytical limit of detection (LOD) assays, dilutions of RNA stock solution were prepared in RNA reconstitution buffer (GeneLink). LAMP primers and guide RNA (gRNA) targeting the N and E genes of SARS-CoV-2 and RNase P gene of human DNA were originally published by Broughton et al(3), and the sequences are listed below in Table S1. LAMP primers (Elim Biosciences) were reconstituted in nuclease free water and gRNAs (IDT) were reconstituted in RNA reconstitution buffer.

For ITP co-focusing visualization experiments, the Mtb target DNA sequence was used (Table S1). 1 μM stock solution of Mtb dsDNA was prepared by pre-hybridizing complementary ssDNA templates (Elim Biosciences) in a buffer containing 50 mM Tris-HCl, 5 mM MgCl_2_, and 1 mM EDTA at 37°C. We designed a Cy5-labeled gRNA (IDT, Table S1) to target the Mtb dsDNA sequence.

### Preparation of contrived samples for ITP-CRISPR detection assay

Forty 60 μL nucleic acid extracts from 400 μL of negative NP swab samples were acquired from the Stanford Clinical Virology Lab with the approval of Stanford University IRB (protocol #48973). The forty extracts were pooled to provide human genomic DNA control from clinical samples. Synthetic SARS-CoV-2 RNA of known concentrations were combined with the pooled clinical nucleic acid extracts before performing analytical LOD experiments for the ITP-CRISPR assay.

### Microfluidic chip and preparation

ITP-based nucleic acid extraction and ITP-CRISPR detection were performed using off-the-shelf glass microfluidic chips (model NS12AZ, Caliper Life Sciences). A single chip consists of two cross geometry channels wet-etched to a 20 μm depth with a 50 μm mask width, resulting in a channel width of 90 um and a roughly D-shaped cross section. The main channel length between the positive/negative electrodes is 72 mm (Fig. S10). To avoid cross-contamination, ensure run-to-run repeatability, and provide uniform surface properties, the channels were rinsed in the following order before each ITP experiment: 10% bleach for 2 min, DI water for 2 min, 1% Triton-X for 2 min, DI water for 2 min, 1 M NaOH for 2 min, and DI water for 2 min. Between each rinse step, the channel was completely dried using vacuum. The buffer loading procedure and buffer placement in the channel sections are detailed in Fig. S10.

### ITP extraction of total nucleic acids

ITP was used to extract total nucleic acids from 10 μL of primary nasopharyngeal (NP) swab clinical samples in viral transport medium. The samples were acquired from the Stanford Clinical Virology Lab with the approval of Stanford University IRB. 10 μL of NP sample was mixed with a 1.1 μL of 10x lysis buffer and incubated at 62°C for 2 min. The 1x composition of lysis buffer included 1.5 % Triton X, 1 mg/mL of Proteinase K, 0.1 mg/mL of carrier RNA (Thermo Fisher). Following incubation, 1 μL of 300 mM HEPES buffer was added, and 10 μL of this mixture was dispensed in the trailing electrolyte (TE) reservoir on-chip (Fig. S10). The leading electrolyte (LE) buffer in the main channel consisted of 100 mM Tris-HCl (pH 7.5), 1 U/μL RNasin Plus, 0.2 % Triton X, 1% of 1.3 MDa Polyvinylpyrrolidone (PVP) and 1x SYBR Green I. SYBR Green I was used to visualize the ITP peak which contained nucleic acids (Fig. 2d). The 10 μL extraction buffer in the LE reservoir consisted of 20 mM Tris-HCl (pH 7.5), 1 U/μL RNasin Plus, and 0.1 mg/mL of carrier RNA. A lower buffer concentration was used for extraction to ensure compatibility with downstream qPCR and LAMP amplification. ITP extraction of nucleic acids was performed at constant voltage of 1 kV supplied by a Keithley 2410 high voltage sourcemeter (Fig. S11).

### RT-LAMP reactions

RT-LAMP reactions were carried out with the WarmStart LAMP Kit (NEB) using the manufacturer’s recommended protocol. The final concentrations of LAMP primers were 1.6 μM for FIP and BIP, 0.2 μM for F3 and B3, and 0.8 μM for LF and LB, as used in Broughton et al (3). Reactions were performed with a final volume of 10 μL, and were set up separately for N, E, and RNase P genes. LAMP reaction mixtures were incubated at 62°C for 20 min.

For ITP-CRISPR analytical LOD experiments, 4 μL of contrived sample containing a mixture of viral RNA (2 μL) and pooled negative clinical NP swab extracts (2 μL) was used as template. For tests involving the complete 30-min assay on clinical patient samples, we used 3 μL of ITP-extracted nucleic acids as template for each LAMP reaction.

### Cas12-gRNA complex preparation

A 10x Cas12-gRNA complex mixture was prepared by pre-incubating 1 μM of LbCas12a (NEB) with 1.25 μM gRNA in 1x NEBuffer 2.1 at 37°C for 30 min. LbCas12-gRNA complexes were prepared independently for N, E, and RNase P genes.

For ITP co-focusing visualization experiment in Fig. 1c, a 10x Cas12-gRNA complex was prepared using 1 μM of LbCas12a (NEB) and 0.5 μM of Cy5-labeled gRNA. Here, a molar excess of LbCas12a was used to minimize free, unbound gRNA.

### ITP-CRISPR detection

The LE buffer consisted of 200 mM Tris, 100 mM HCl, 10 mM MgCl_2_, and 0.1 % PVP. The TE buffer consisted of 100 mM Tris, 50 mM HEPES, 10 mM MgCl2, 0.1 % PVP and 250 nM of ssDNA fluorescence-quencher reporter (/56-FAM/TTATT/3IABkFQ/, IDT). Before each SARS-CoV-2 ITP-CRISPR detection experiment, 2 μL of the 10x LbCas12-gRNA complex was combined with 2 μL of the corresponding LAMP amplicon and 16 μL LE buffer. For ITP co-focusing visualization experiments in Fig. 1c, 2 μL of the 10x LbCas12-gRNA complex was combined with 2 μL of pre-prepared Mtb dsDNA template and 16 μL LE buffer. The on-chip buffer loading procedure is described in Fig. S10.

The ITP-CRISPR detection experiments were performed at constant current of 4 μA supplied by a Keithley 2410 sourcemeter (Fig. S11). Fluorescence images of the moving ITP peak were acquired in 30 s intervals using a CMOS camera (Hamamatsu ORCA-Flash4.0) mounted on an inverted epifluorescence microscope (Nikon Eclipse TE200). For widefield images of ITP peak in Fig. 1b, we used a microscope objective (Nikon) with 2x magnification and 0.1 NA objective to enable imaging over a wide field of view. For all other quantitative fluorescence measurements of the ITP peak, a 10x magnification and 0.4 NA (Nikon) objective was used.

For ITP co-focusing visualization experiments of Fig. 1c, a white LED (Thorlabs) excitation source was used to enable simultaneous imaging of a Cy5-labeled (red channel) gRNA of the Cas12-gRNA complex and cleaved FAM-labeled (green channel) ssDNA reporter molecules. During the experiment, we manually switched between filter cubes for the green and far-red emission wavelengths. For all other experiments involving ITP-based nucleic acid extraction and ITP-CRISPR assay quantification, we used blue LED excitation source with a green emission filter cube.

### Image analysis of fluorescence readouts

The fluorescence signal was calculated from raw experimental images using ImageJ software (National Institutes of Health). Fluorescence intensity values were integrated over a pre-defined square region around the ITP peak. The dimension of the square region was around 4 channel widths. A background value was obtained by integrating the signal over a square region with same dimensions in the same image and in a region significantly away from the ITP peak. The reported signal is the background subtracted integrated fluorescence intensity.

### RT-qPCR assay

The RT-qPCR assay was performed using the ABI 7500 Fast DX (Applied Biosystems) instrument. We performed assays for the E and RNase P genes separately in 20 μL reaction volumes using the Luna Universal Probe One-Step RT-qPCR Kit (New England Biolabs). The final concentrations of primer and probe were 400 and 200 nM, respectively. We followed the recommended protocol in Corman et al.(1) For quantification from clinical samples, we used 8 and 2 μL of ITP-extracted nucleic acids for the E gene and RNase P gene reactions, respectively. For the E gene standard curve, we used 5 μL of various dilutions of synthetic RNA controls as template.

### Human clinical sample collection and preparation

Clinical positive and pooled negative SARS-CoV-2 nasopharyngeal (NP) swab samples in viral transport medium (VTM) were collected at the Stanford Clinical Virology lab with the approval of Stanford University IRB. For quantification experiments, the original positive clinical nasopharyngeal swab specimen (Covid19-P1) was diluted 1:10 (Covid19-P2), 1:100 (Covid19-P3), 1:1000 (Covid19-P4) in VTM from pooled negative clinical NP swab specimens. The negative samples (Covid19-N1, Covid19-N2, Covid19-N3, Covid19-N4) were four aliquots of the pooled negative clinical NP swab sample.

## Acknowledgements

A.R. gratefully acknowledges support from the Bio-X Bowes Fellowship of Stanford University. D.A.H. is supported by a National Science Foundation Graduate Research Fellowship. J.G.S., A.R., D.A.H., and N.B. acknowledge funding from the Stanford Chemistry, Engineering & Medicine for Human Health (ChEM H) institute.

## Author contributions

A.R. and J.G.S. conceived of the assay design. A.R. developed detailed protocols, designed and carried out experiments. A.R and D.A.H. processed experimental data and performed the analysis. D.A.H., A.R. and J.G.S. designed the figures. B.A.P. and M.K.S. provided clinical samples and advice regarding traditional techniques. E.S. helped with an initial design of experiments. J.G.S. supervised the project, and N.B. and B.A.P. helped supervise the project. All authors provided critical feedback that helped shape the research, analysis and manuscript.

## SI Appendix

**Figure S1.**
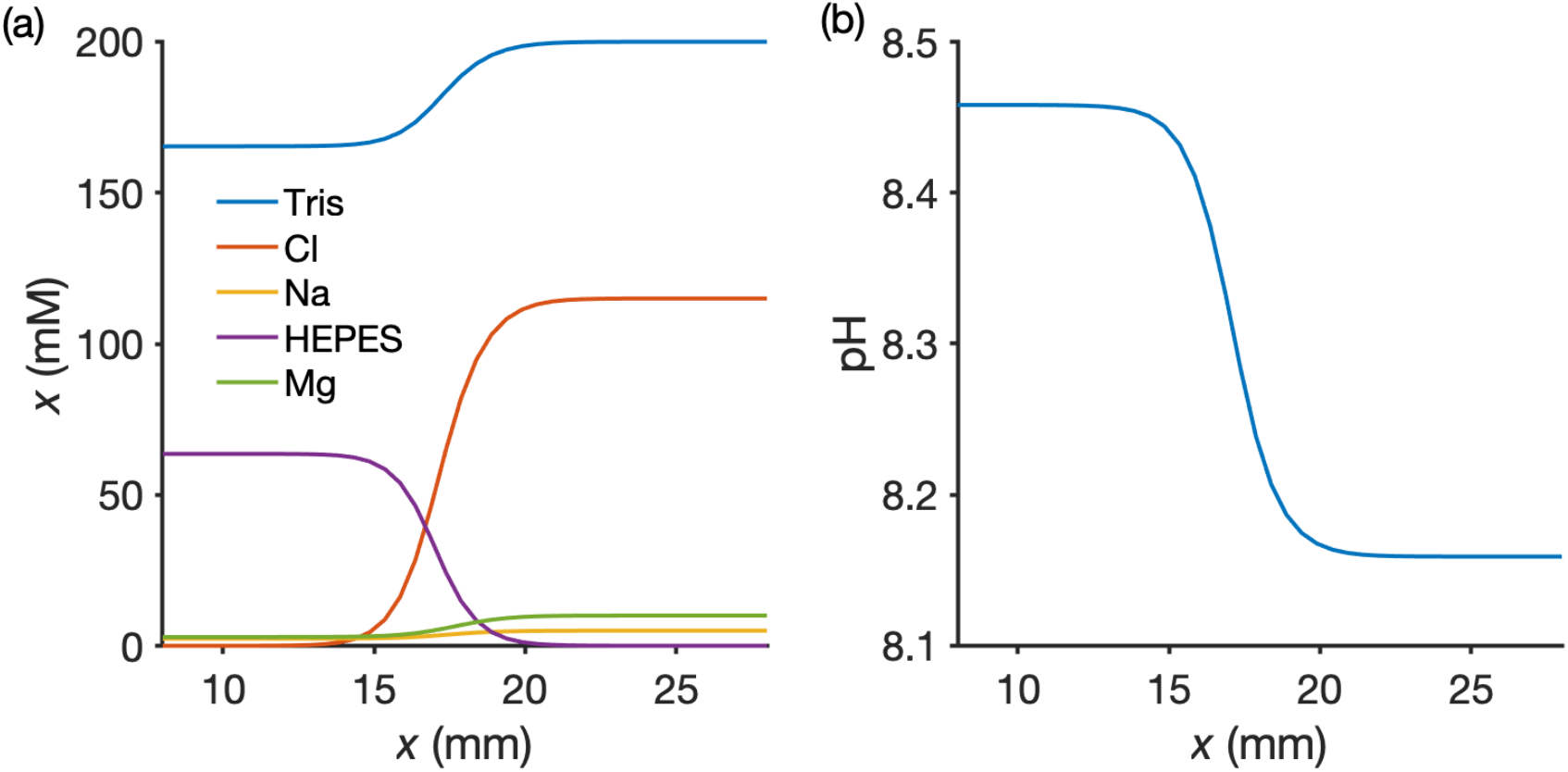
Simulation of ITP-CRISPR detection using Spresso. (12). (a) Concentration of ions versus *x* near the LE-TE interface. Applied electric field is in the direction from right to left. Not shown is the concentration of analyte ions (focused at the LE-TE interface region) since they are several orders of magnitude smaller than the buffer ions. (b) pH versus *x* near the ITP peak region. Note that the pH in the LE-TE interface varies between 8.16 (in the LE region) and 8.46 (in the adjusted TE region).

**Figure S2.**
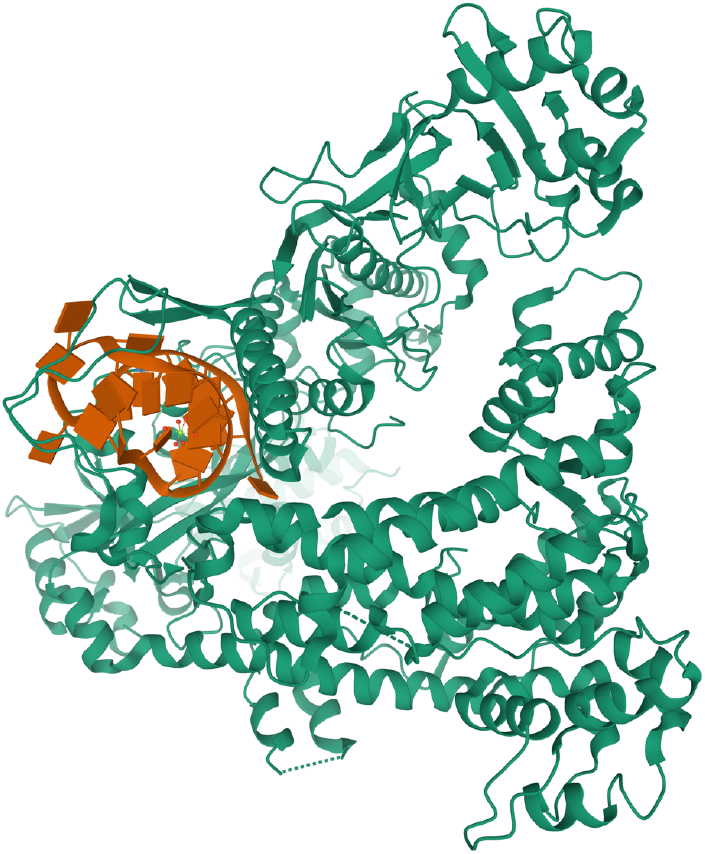
Structure of Cpf1(Cas12a)/RNA complex generated using PyMOL (PDB: 5id6) (13) and calculation of theoretical isoelectric point. The source organism is wild-type Lachnospiraceae bacterium ND2006 (same as that of EnGen LbaCas12a, NEB). The theoretical isoelectric point (pI) computed using the ExPASy tool (compute pI/Mw) for this entire LbaCas12a protein sequence (UniProtKB: A0A182DWE3) is 8.39. When Cas12 (shown in green) complexed with RNA (shown in orange) is present in a buffer of pH around ~8.3 (between LE and TE), we hypothesize that the combined Cas12a-gRNA complex has a net negative charge in solution. Further, based on experiments and simulations, we estimate the effective electrophoretic mobility of Cas12a-gRNA complex is between −2.09 x 10^−8^ m^2^/V-s (HEPES) and −7.91 x 10^−8^ m^2^/V-s (Chloride) for our ITP chemistry.

**Figure S3.**
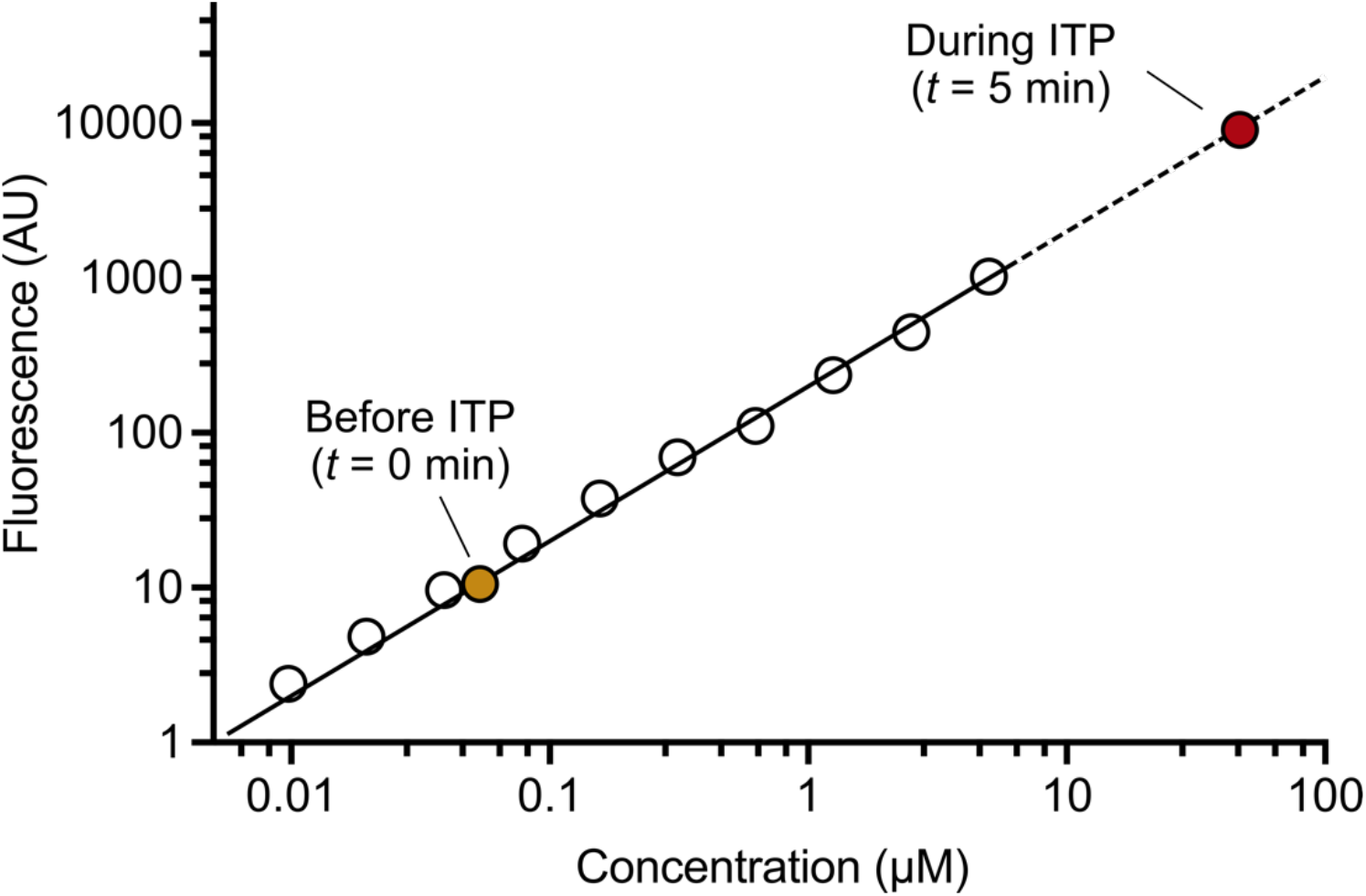
Fluorescence calibration curve. Various concentrations of Cy5-tagged gRNA (open symbols) were used to generate a fluorescence calibration curve. ITP pre-concentrates Cas12-gRNA, reporters, and target DNA by order ~1000-fold compared to initial loading concentrations. All these molecules are concentrated from ~0.1 μL initial volume (on-chip, before ITP) to a ~100 pL volume (on-chip, during ITP). The increase in concentration of enzyme and substrate can significantly accelerate production rates.

**Figure S4.**
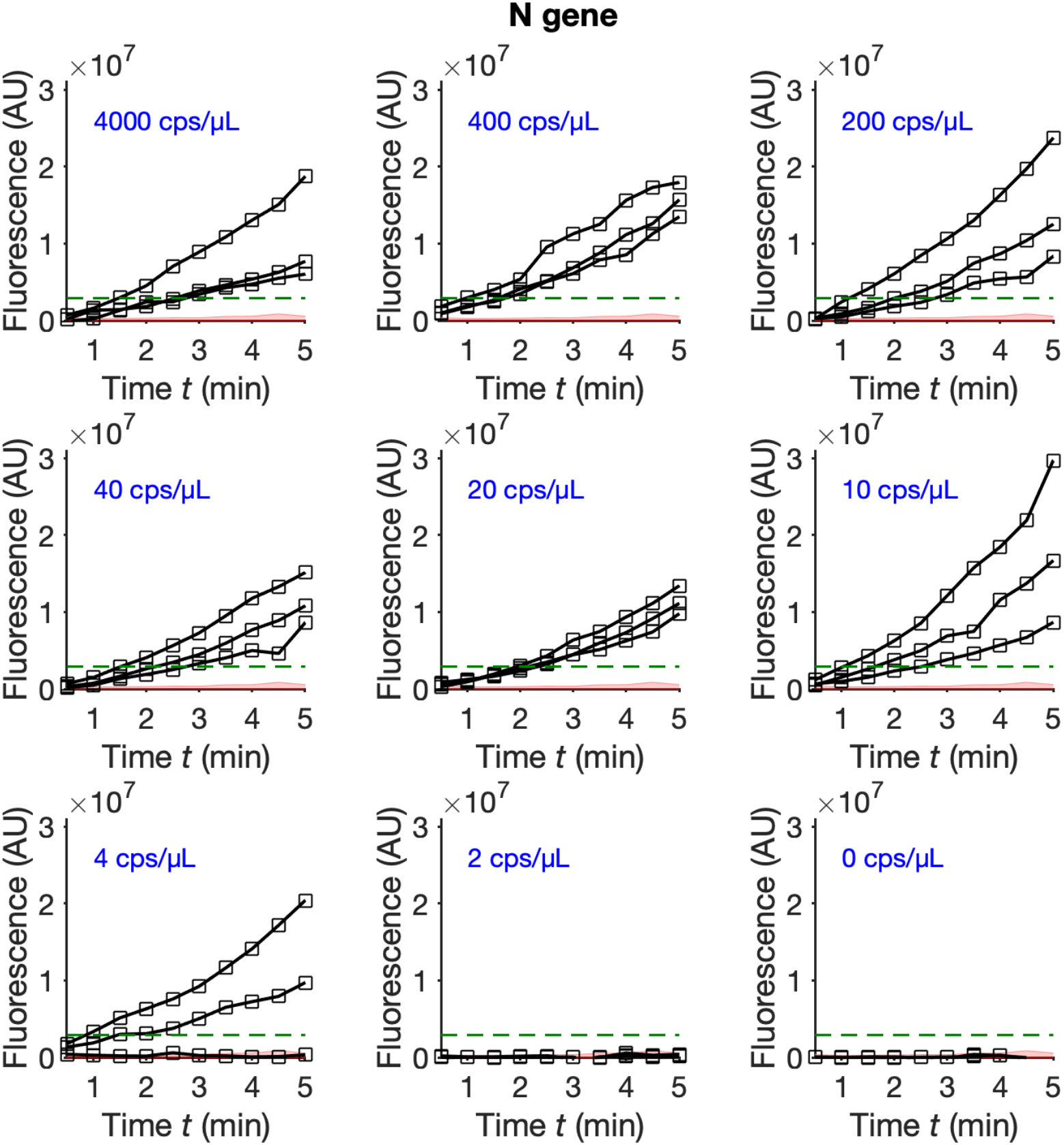
Experimental measurements of fluorescence signal for ITP-CRISPR detection of N gene in contrived samples. Area-integrated image fluorescence intensity (in the microfluidic channel) versus time during N gene ITP-CRISPR detection on contrived samples. Data was recorded every 30 s. Three replicates are shown for each concentration. Dashed lines represent the threshold signal for the assay.

**Figure S5.**
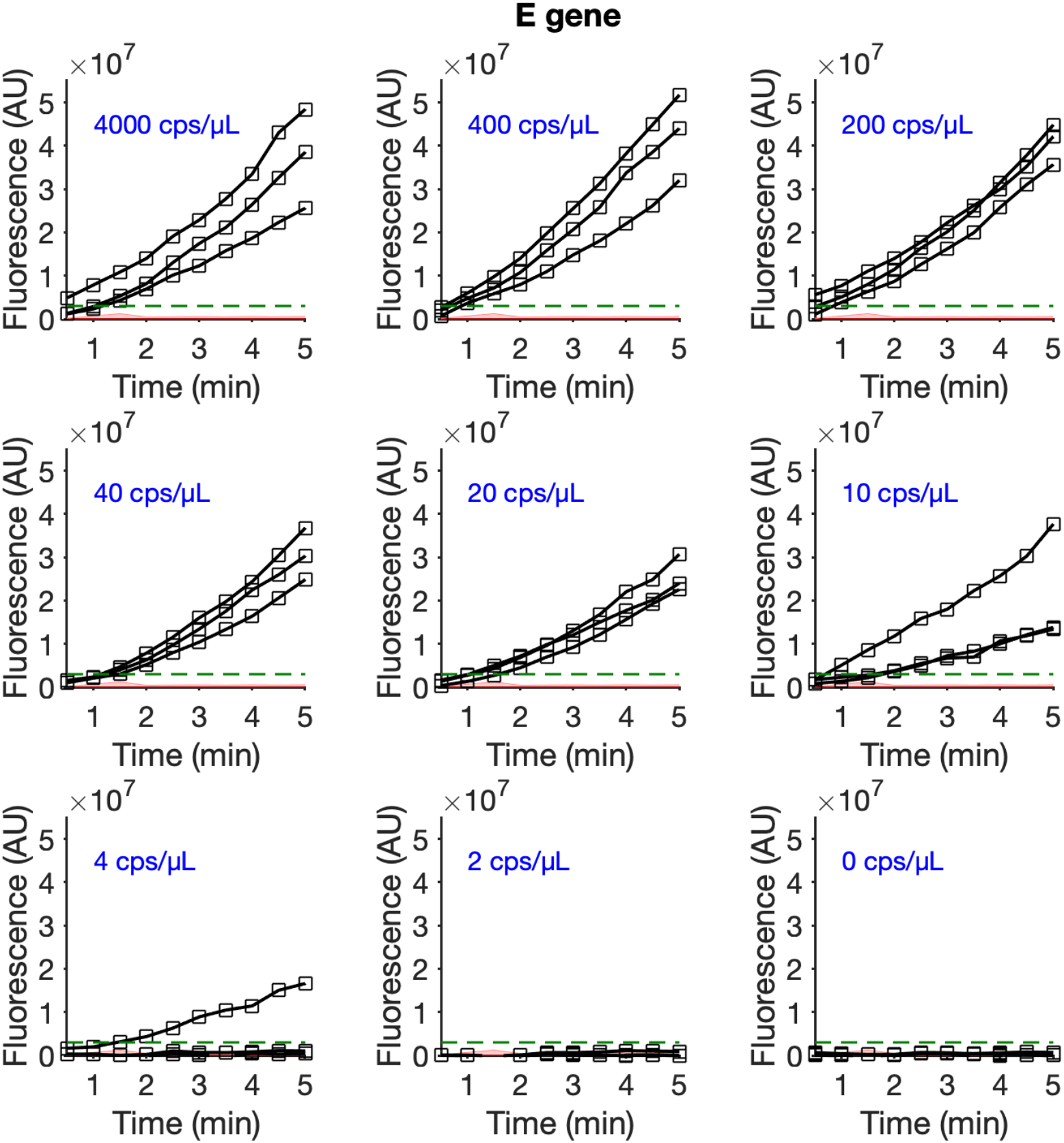
Experimental measurements of fluorescence signal for ITP-CRISPR detection of E gene in contrived samples. Area-integrated image fluorescence intensity (in the microfluidic channel) versus time during E gene ITP-CRISPR detection on contrived samples. Data was recorded every 30 s. Three replicates are shown for each concentration. Dashed lines represent the threshold signal for the assay.

**Figure S6.**
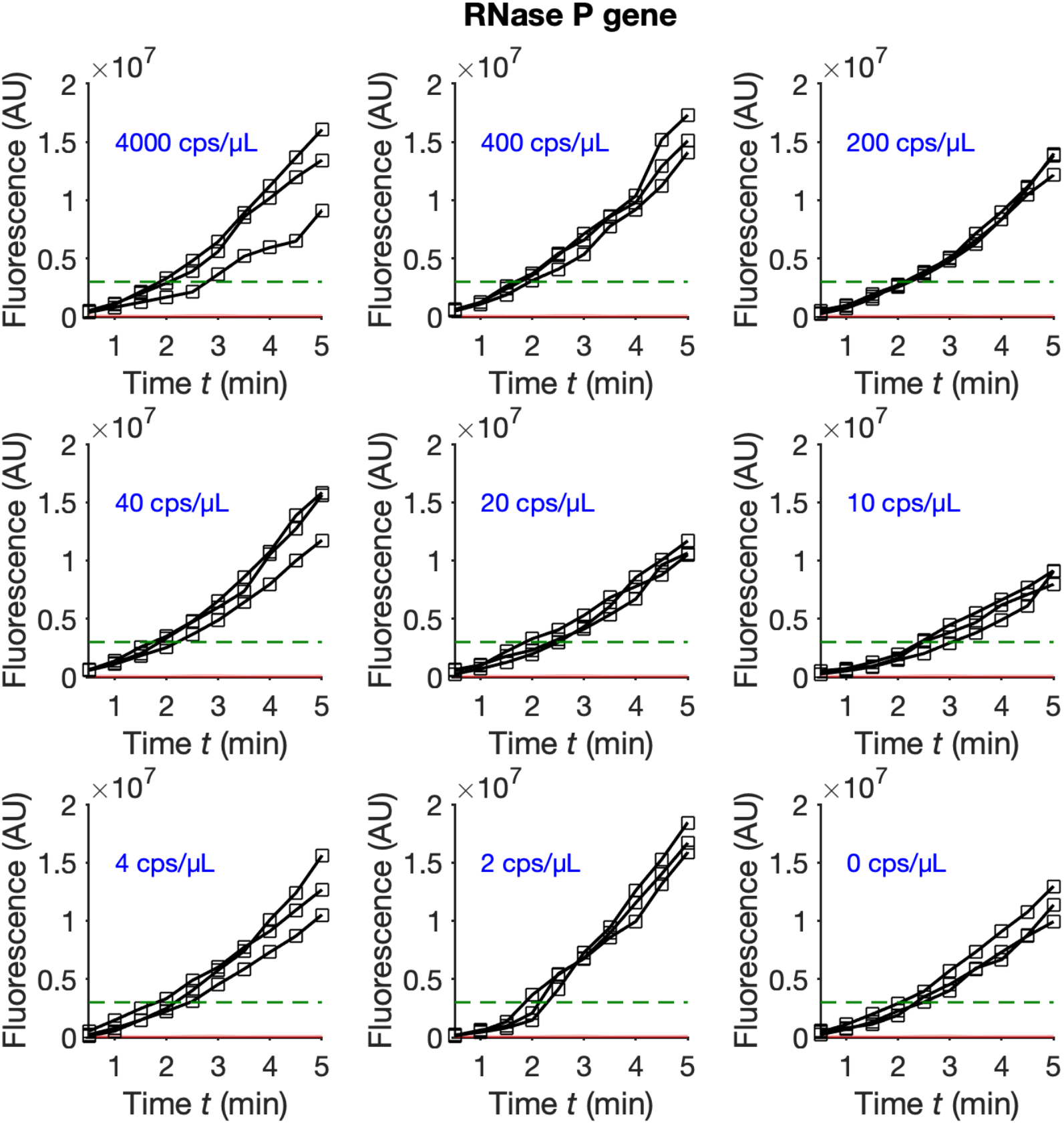
Experimental measurements of fluorescence signal for ITP-CRISPR detection of RNase P gene in contrived samples. Area-integrated image fluorescence intensity (in the microfluidic channel) versus time during RNase P gene ITP-CRISPR detection on contrived samples. Data was recorded every 30 s. Three replicates are shown for each concentration. Dashed lines represent the threshold signal for the assay. Note that human DNA from pooled negative nasopharyngeal swab extracts was present as background in all contrived samples.

**Figure S7.**
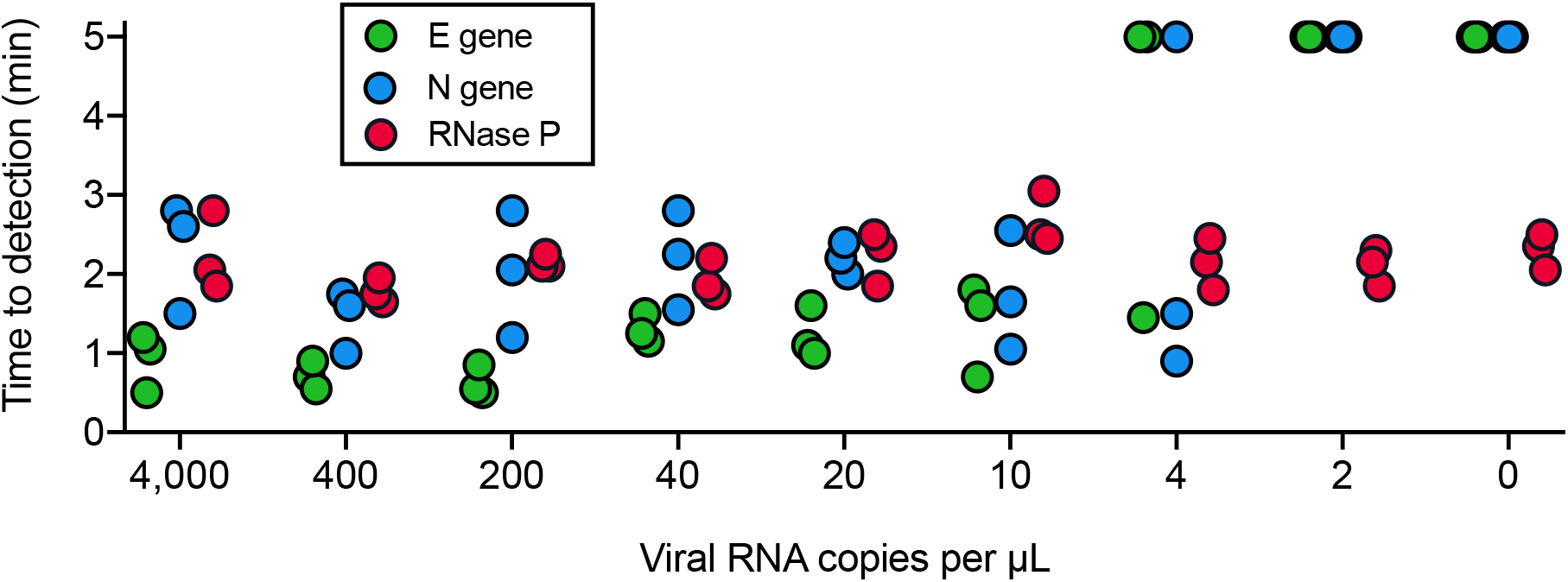
Time to signal detection on contrived samples for the ITP-CRISPR assay. A positive detection occurs when the measured fluorescence signal crosses a threshold value. The time to signal detection is the time at which the measured signal crossed the threshold value. Samples for which signal did not exceed the threshold after 5 min of assay time are denoted by symbols at 5 min.

**Figure S8.**
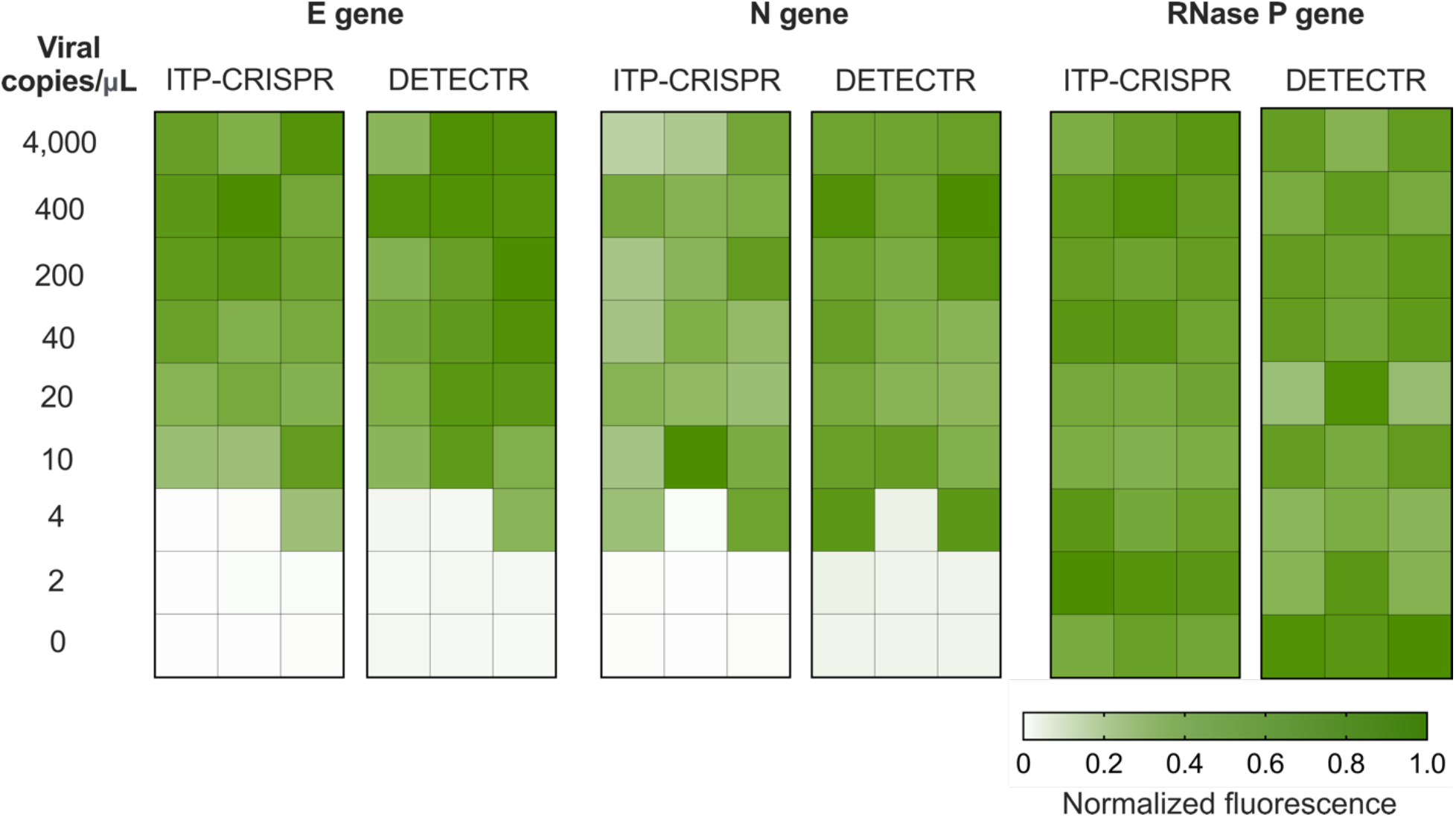
Comparison of ITP-CRISPR and DETECTR-based assays for SARS-CoV-2 detection on contrived samples. Shown are normalized fluorescence values for three replicates of ITP-CRISPR and DETECTR-based detection of N gene, E gene, and RNase P genes on contrived samples. For each method and target gene, fluorescence is normalized to the maximum value across all samples. For the DETECTR method, we followed the protocol in Broughton et al(3) and used endpoint fluorescence readout at 10 min.

**Figure S9.**
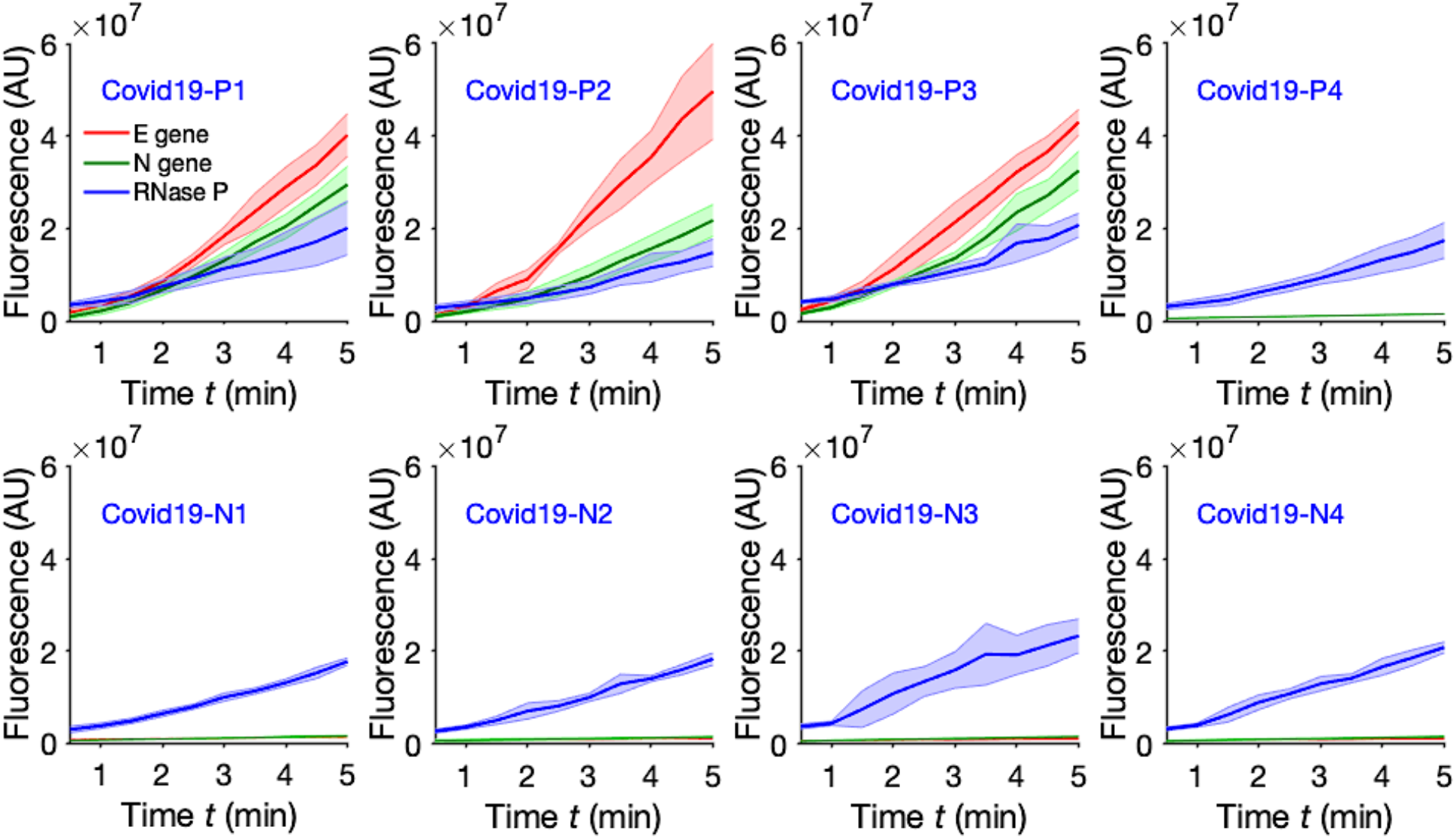
Measured kinetic curves for ITP-CRISPR detection of N gene, E gene, and RNase P gene on clinical samples. Four positive (Covid19-P1 to Covid19-P4) and four negative (Covid19-P1 to Covid19-P4) nasopharyngeal swab samples were each tested in three technical replicates (*n* = 3). Fluorescence signal was measured every 30 s. Solid lines represent the mean signal and the shaded envelope represents the standard deviation. Covid19-P4 was below the LOD of our assay as confirmed by qPCR (Fig. 2e).

**Figure S10.**
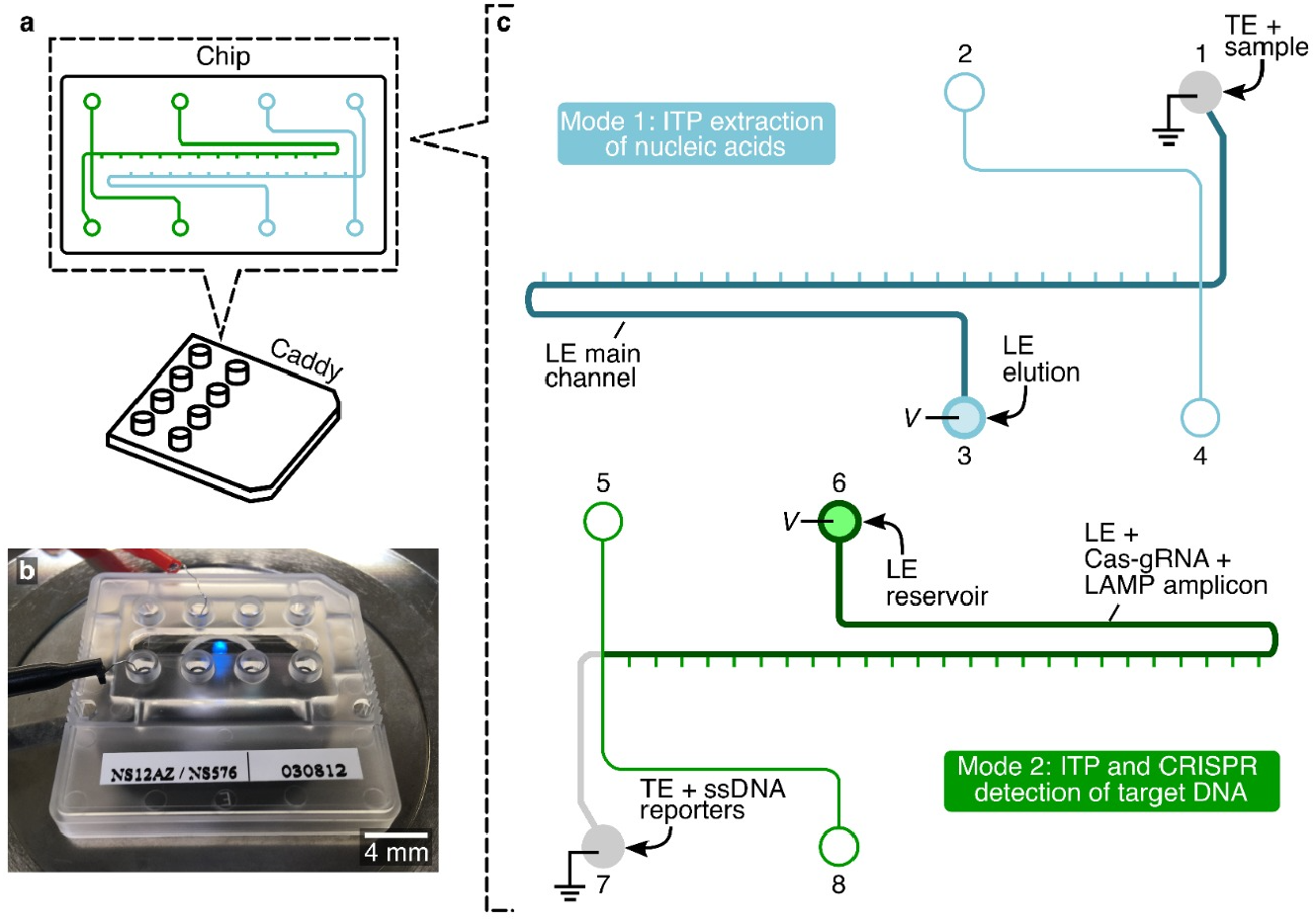
Experimental setup and microfluidic chip layout. (a) Schematic of the glass chip bonded to a plastic caddy using epoxy. The chip consists of two identical cross-channels adjacent to each other. (b) Experimental setup showing the chip and electrodes on a microscope stage during Mode 2 operation (c) Buffer placement and sample loading in Mode 1 and Mode 2. In Mode 1, first, reservoirs 2, 3, and 4 are filled with 10 μL of LE main channel buffer and vacuum is applied at reservoir 1 till the main channel is completely filled with LE. Then, reservoirs 3 and 1 are emptied and filled with 10 μL of LE elution buffer and lysed sample in TE, respectively. A constant voltage of 1 kV is applied for ~3 min. For Mode 2, first, 5 μL of LE combined with Cas12-gRNA and LAMP amplicon is loaded in reservoir 6, 5 μL of LE is loaded in reservoir 8, and 5 μL TE combined with ssDNA reporters is loaded in reservoir 7. Vacuum is applied at reservoir 5 briefly till the channels are filled as depicted in the schematic. Then, reservoirs 5, 6, and 8 are emptied and loaded with 10 μL of LE, and reservoir 7 is emptied and loaded with 10 μL TE. A constant current of 4 μA is applied for 5 min and fluorescence intensity of the ITP peak is recorded using a CMOS camera every 30 s.

**Figure S11.**
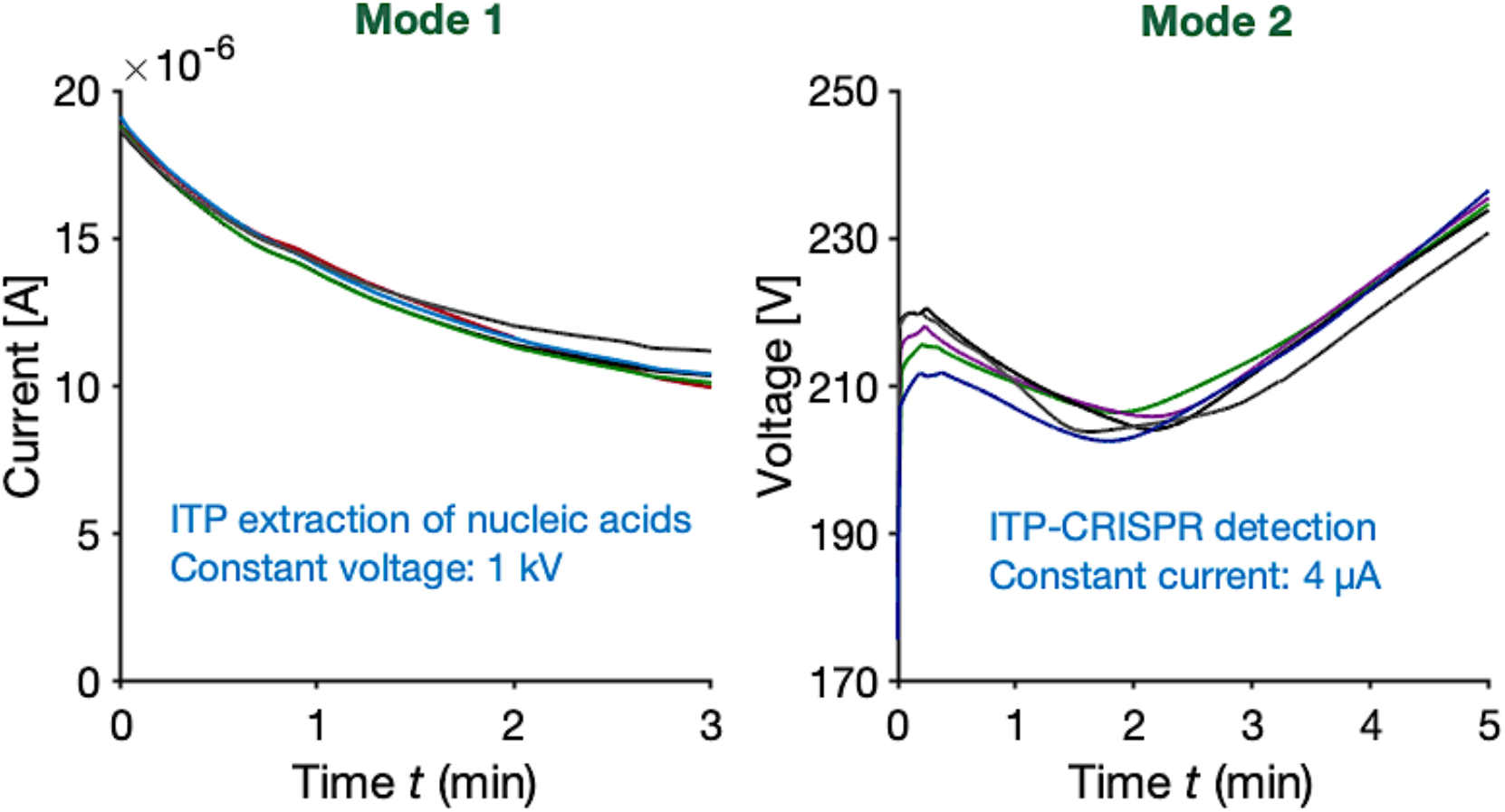
Measured current and voltage versus time during ITP extraction (stage 1) and ITP-CRISPR detection (stage 2). In operational Mode 1, a constant voltage of 1 kV was applied for 3 min to complete ITP extraction of nucleic acids from nasopharyngeal swab sample. In Mode 2, a constant current of 4 μA was applied for 5 min for ITP-CRISPR-based detection. Shown are current and voltage traces from five typical experiments to demonstrate the reproducibility of the process.

**Figure S12.**
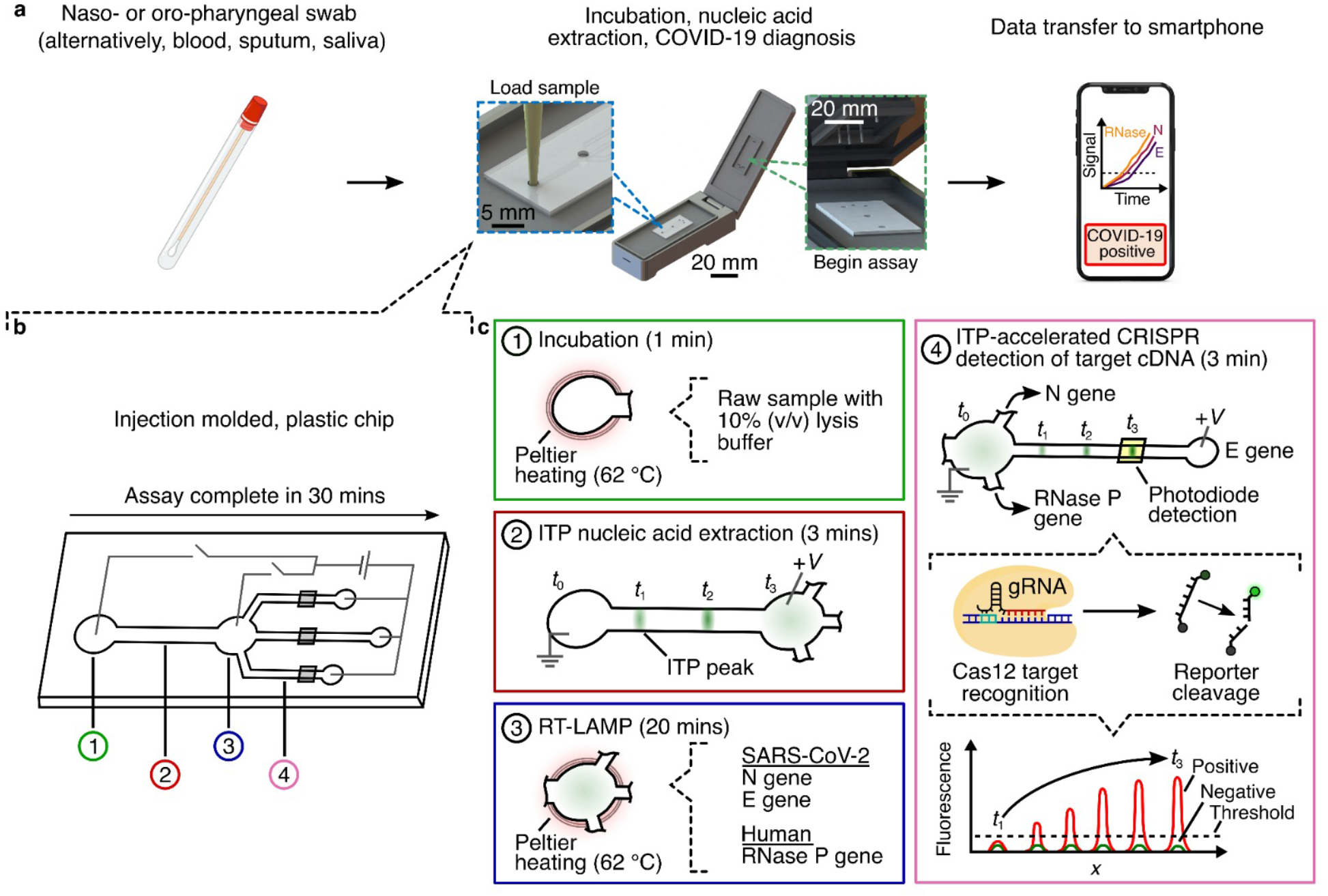
Schematic for a fully integrated chip design on a portable device. An injection molded plastic chip fits into a hand-held portable device powered and controlled via USB includes laser-induced fluorescence detector, DC-to-DC converters, a photodiode, and integrated microheaters (for LAMP incubation at 62°C). See Bercovici et al(14) for an integration of at least one ITP assay into a portable device. We propose here the concept that such a system can integrate ITP-based nucleic acid extraction, multiplexed isothermal amplification of target cDNA of N and E genes of SARS-CoV-2 and RNase P control, followed by ITP-CRISPR-based cDNA detection in three separate channels using photodiodes.

**Table S1.**
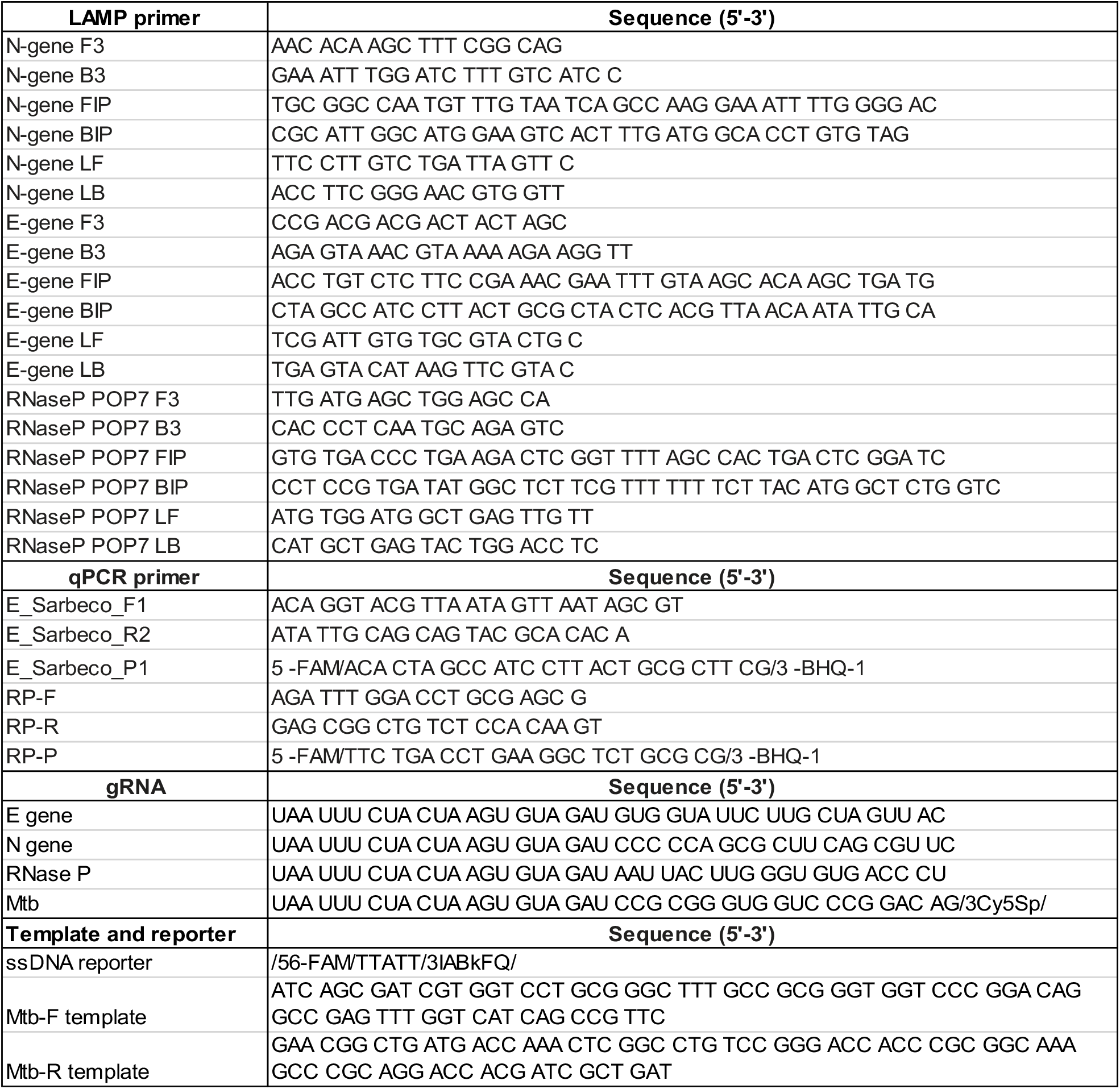
List of gRNAs, LAMP primers, RT-qPCR primers, template and reporter sequences. Mtb sequences were used for ITP co-focusing experiments of Figure 1c. E gene, N gene, and RNase P sequences (originally published in Broughton et al (3)) were used for SARS-CoV-2 detection. The ssDNA reporter was used commonly for all ITP-CRISPR experiments.

